# A dendritic mechanism for balancing synaptic flexibility and stability

**DOI:** 10.1101/2022.02.02.478840

**Authors:** Courtney E. Yaeger, Dimitra Vardalaki, Qinrong Zhang, Trang L.D. Pham, Norma J. Brown, Na Ji, Mark T. Harnett

**Affiliations:** McGovern Institute for Brain Research, MIT, Cambridge, MA, USA; Department of Brain and Cognitive Sciences, MIT, Cambridge, MA, USA; Department of Physics, University of California, Berkeley, CA, USA; Department of Molecular and Cell Biology, University of California, Berkeley, CA, USA; Helen Wills Neuroscience Institute, University of California, Berkeley, CA, USA; Molecular Biophysics and Integrated Bioimaging Division, Lawrence Berkeley National Laboratory, Berkeley, CA, USA

## Abstract

Biological and artificial neural networks learn by modifying synaptic weights, but it is unclear how these systems can retain previous knowledge and also acquire new information. Here we show that cortical pyramidal neurons can solve this plasticity-versus-stability dilemma by differentially regulating synaptic plasticity at distinct dendritic compartments. Oblique dendrites of adult mouse layer 5b cortical pyramidal neurons selectively received monosynaptic thalamic input, integrated linearly, and— surprisingly—lacked synaptic potentiation. In contrast, basal dendrites, which do not receive thalamic input, exhibited conventional NMDA receptor-mediated supralinear integration and synaptic potentiation. Congruently, spiny synapses on oblique branches showed low structural plasticity in vivo. A selective decline in NMDAR activity and expression at synapses on oblique dendrites was controlled by an experience-dependent critical period. Our results demonstrate a new biological mechanism for how single neurons can safeguard a set of inputs from ongoing plasticity by altering synaptic properties at distinct dendritic domains.

## Main

The brain learns continuously by balancing plasticity and stability, whereby new information is incorporated without disrupting prior knowledge. Synaptic plasticity is the key principle for learning, but neural circuits must also exhibit sufficient synaptic stability to maintain information storage over the long term. This process poses a major unsolved problem for both biological and artificial neural networks, known as the plasticity-stability dilemma^1–4^. In artificial learning systems, this problem manifests as catastrophic forgetting, where previously acquired information is overwritten unless weight changes are limited^5–7^. While half a century of experimental and theoretical work on biological synaptic plasticity has focused on how plasticity is induced^8–17^ and maintained within a working range^18–22^, little is currently known about how neurons sustain previously learned information throughout a lifetime of experience-dependent plasticity.

In the mammalian sensory cortex, a classical plasticity-stability challenge occurs during early postnatal development^23^. Initial sensory experiences must be stored and stabilized for robust sensory processing, but cortical circuits must maintain flexibility to accommodate subsequent learning. For example, primary sensory inputs in the visual cortex (V1) from the dorsal lateral geniculate nucleus of the thalamus (dLGN) rapidly change at the onset of vision^24–27^. By adulthood, these inputs are stable and no longer plastic during normal visual experience^28–31^. However, higher-order cortical representations remain highly plastic^32–35^. Because single cortical neurons receive spatially-targeted inputs^36–39^, we hypothesized that dendritic compartmentalization could arbitrate plasticity or stability for different kinds of information. Using complimentary physiological and imaging techniques, we identified dLGN-recipient and non-recipient dendritic domains within layer 5b pyramidal neurons (L5b PNs) and characterized the synaptic properties and plasticity of these domains across maturation.

## Results

### L5b PN oblique dendrites selectively receive thalamic input and exhibit distinct integrative properties from basal dendrites

Visual information from dLGN projects primarily to cortical layer 4^40^. First-order thalamic inputs are known to synapse on L5 PNs^41–43^, which integrate inputs from all cortical layers, though the location of the synaptic contacts is unclear. Physiological data from juvenile V1^38^ and anatomical data^44^ suggest that dLGN inputs target the apical oblique dendrites of L5 PNs, which can reside in layer 4. To directly test where dLGN inputs make functional synapses on L5b PNs in adult mouse V1, we employed subcellular channelrhodopsin (ChR2)-assisted circuit mapping^39^. After viral expression of ChR2 in dLGN (Fig. 1a), we performed whole-cell patch-clamp recordings from V1 L5b PNs in acute slices from adult mice (postnatal day 56+). Local photostimulation of ChR2-expressing dLGN boutons across the dendritic tree revealed monosynaptic connections tightly restricted to apical oblique branches (Fig. 1b,c).

**Figure 1:**
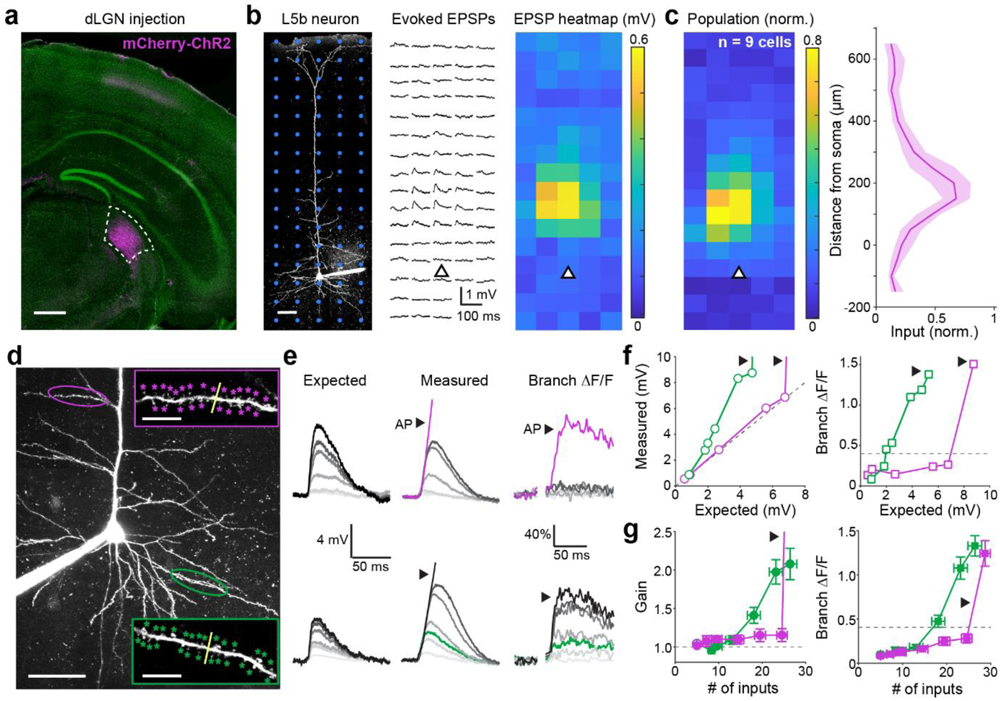
Apical oblique dendrites of adult V1 L5b PNs selectively receive dLGN input and do not have NMDAR-mediated integrative properties. a. Example confocal image of a coronal brain slice showing dLGN neurons infected with AAV2-hSyn-mCherry-ChR2. Scale bar: 500 µm. b. Left: Example two-photon image of a V1 L5b PN whole-cell recording with overlaid photostimulation grid (blue). Scale bar, 50 µm. Middle: Average evoked EPSPs from 6 repetitions of 473 nm photostimulation for the neuron and grid shown. Soma position is indicated by the triangle. Right: Heatmap of average evoked EPSPs. Each square is 50 µm. c. Left: Normalized heatmap of pooled L5b PNs (n = 9 neurons, 9 slices, 3 P56+ animals), aligned at the soma. Each square is 50 µm. Right: Mean normalized input as a function of depth for all neurons. Shaded region indicates the bootstrapped 95% confidence interval. d. Example two-photon z-stack of a L5b PN with glutamate uncaging sites indicated on an oblique (purple) and a basal (green) branch, scale bar: 50 µm. Uncaging sites (stars) and linescan (yellow) are magnified in insets (scale bars: 10 µm). e. Expected EPSPs, measured EPSPs, and local branch Ca^2+^ signals for oblique (top) and basal (bottom) dendrites shown in d. Left: Expected voltages from increasing numbers of linearly summed inputs. Middle: Measured EPSPs in response to increasing numbers of synchronous uncaging inputs. Stimulation of all inputs led to an action potential (AP, arrow). Right: Corresponding changes in local branch OGB-1 signal (ΔF/F). f. Measured voltage (left) and branch ΔF/F (right) as a function of expected voltage for the branches shown in d. Dashed line indicates linearity (left) and threshold Ca^2+^ signal (right). g. Adult population gain (measured/expected, left) and branch ΔF/F (right). n = 13 basal branches from 7 animals and 9 oblique branches from 6 animals. All animals were P56+. Basal APs removed for clarity. Basal vs oblique gain (AP excluded), Mann-Whitney U test: p = 7.63E-04; basal vs oblique ΔF/F (AP excluded), Mann-Whitney U test: p = 1.08E-05.

In light of this specific targeting of dLGN inputs to L5b PN oblique dendrites, we asked whether these branches exhibited distinct properties compared to nearby basal dendrites (Fig. 1d). In acute slices from adult mice, basal dendrites displayed highly supralinear integration and large local Ca^2+^ signals in response to spatiotemporally-clustered two-photon glutamate uncaging (Fig. 1e,f,g), consistent with previous reports of NMDA spikes in juvenile cortex^45^. In contrast, oblique dendrites integrated inputs linearly until axosomatic action potential threshold was reached and did not show significant subthreshold Ca^2+^ influx (Fig. 1e,f,g), similar to what has been previously described in retrosplenial cortex^36^. These differences could not be explained by the size of uncaging events, the number of inputs, distance from the soma, action potential threshold, or δV/δt prior to action potential initiation (Extended Data Fig. 1). Thus, oblique dendrites of adult V1 L5b PNs selectively receive dLGN input and exhibit a specialized linear integration mode.

### Oblique dendrites lack synaptic potentiation in adult V1 L5b PNs

The lack of large NMDAR-mediated subthreshold Ca^2+^ signals in oblique dendrites led us to hypothesize that synapses on these branches may exhibit plasticity differences compared to basal dendrites. We tested synaptic potentiation at basal or oblique branches by pairing focal electrical stimulation of presynaptic axons with somatic current injection. A theta glass bipolar stimulating electrode was positioned within 10 µm of either a basal or an oblique dendritic branch (Fig. 2a,b, Extended Data Fig. 2) to generate a 1 to 2 mV excitatory postsynaptic potential (EPSP) in the presence of a GABA-A antagonist. EPSPs were paired with 20 ms somatic current injections that drove bursts of 2-4 action potentials. The peak of the EPSP preceded the peak of the first action potential by 10 ms. 5 pairings at 10 Hz were repeated 30 times, with 10 seconds between epochs (Fig. 2c). This protocol robustly potentiated synapses on basal dendrites (Fig. 2d,f,g, Extended Data Fig. 2), similar to findings in juvenile cortex^12^. Potentiation at basal branches required pre-and post-synaptic coincidence as well as NMDARs (Fig. 2g, Extended Data Fig. 2). In contrast, synapses on oblique dendrites did not exhibit any significant potentiation after pairing (Fig. 2e-g, Extended Data Fig. 2). Doubling the number of pairings did not change this outcome (Extended Data Fig. 2). These findings indicate that, unlike basal dendrites, oblique branches of L5b PNs in adult V1 lack the capability for synaptic potentiation under conventional plasticity induction protocols.

**Figure 2:**
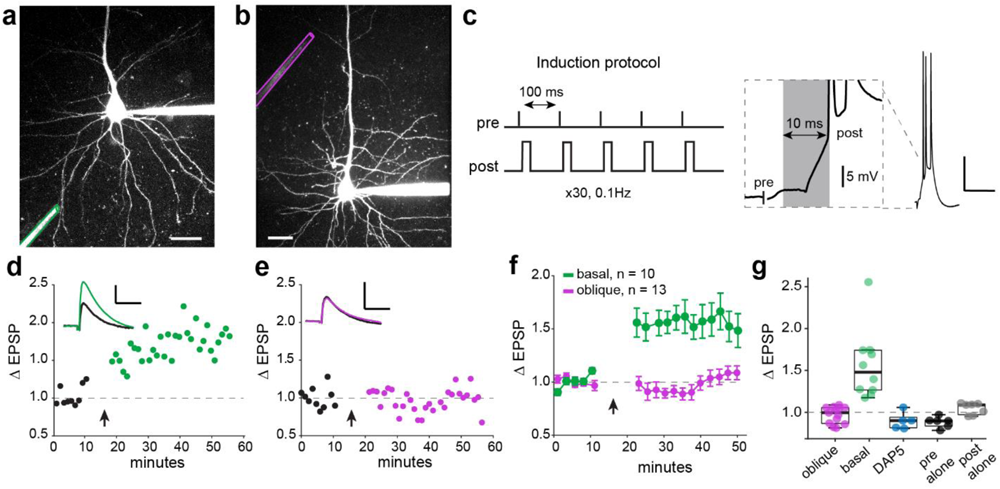
Oblique dendrites in adult V1 L5b PNs lack long-term synaptic potentiation, unlike basal dendrites. a. Example two-photon z-stack of whole-cell recording with focal stimulation (green) near a basal dendrite. Scale bar: 25 µm. b. As in a, but for an oblique dendrite (purple). c. Plasticity induction protocol. 5 pairings of pre-and post-synaptic stimulation at 100 Hz, repeated 30 times at 0.1 Hz. Right: a 10 ms interval between the peak of the evoked EPSP and the peak of the first action potential. Scale bar: 20 mV, 50 ms; inset: 5 mV, 5 ms. d. Example recording from the basal dendrite shown in a. After 10 minutes of baseline recording, the induction protocol is delivered (arrow). Inset shows average EPSP before (black) and after (color) pairing. Scale bar: 1 mV, 25 ms. Responses are 1 min binned averages and normalized to baseline. e. As in d, for the oblique dendrite shown in b. f. Normalized EPSP amplitude for neurons stimulated at basal (green) or oblique (purple) branches (2 min bins; n = 10 basal branches, 10 animals and 13 oblique branches, 12 animals. All animals were P56+.). Error bars are s.e.m. Basal vs. oblique post-induction EPSP, Mann-Whitney U test: p = 4.95E-63. g. Change in EPSP after induction for all oblique branches, basal branches, and basal branches with DAP5 or with only presynaptic or postsynaptic activation during induction. Basal vs. oblique average change in EPSP, Mann-Whitney U test: p = 6.33E-05. All other comparisons not significant. Boxplot: median, 25^th^, and 75^th^ percentiles with +/–2.7σ whiskers.

### Synaptic plasticity in oblique dendrites is limited to a postnatal critical period

A developmental decline in synaptic potentiation has been reported for some cortical synapses ^46,47^, and it is known that dLGN inputs in V1 undergo substantial refinement during early postnatal development^25,48,49^. Therefore, we hypothesized that synapses on oblique L5b PN dendrites may exhibit NMDAR-mediated supralinear integration and plasticity during early postnatal development. We found that at eye-opening (postnatal days 12-14), oblique and basal dendrites displayed similar NMDAR-dependent supralinearities and large local Ca^2+^ signals (Fig. 3a-c, Extended Data Fig. 3). These properties persisted into the canonical critical period (P18-22)^31,50^. But by 4 weeks of age (P28-P32), oblique dendrites had developed adult-like properties, with linear synaptic integration and minimal local Ca^2+^ signals (Fig 3c, Extended Data Fig. 3). Synaptic plasticity at oblique dendrites followed the same developmental trajectory: oblique and basal dendrites exhibited NMDAR-dependent synaptic potentiation at juvenile ages (Fig. 3d, Extended Data Fig. 3) and by P28 apical integrative properties became adult-like. These findings suggested that visual experience mediates the maturation of oblique dendrite properties: to test this, we conducted these same experiments in mice deprived of light from birth. In P28 dark-reared animals, oblique dendrites retained immature supralinear integration with large local Ca^2+^ signals and synaptic potentiation (Fig. 3e, Extended Data Fig. 3). Furthermore, dark-reared mice re-exposed to a normal light cycle for 2-4 weeks did not exhibit adult-like integrative properties (Extended Data Fig. 3), indicating that the oblique compartment matures only during a specific window in development. Collectively, these results reveal a restricted developmental window in which visually-driven thalamic activity shifts the L5b PN oblique dendritic compartment to a linearly-integrating, non-potentiating mode.

**Figure 3:**
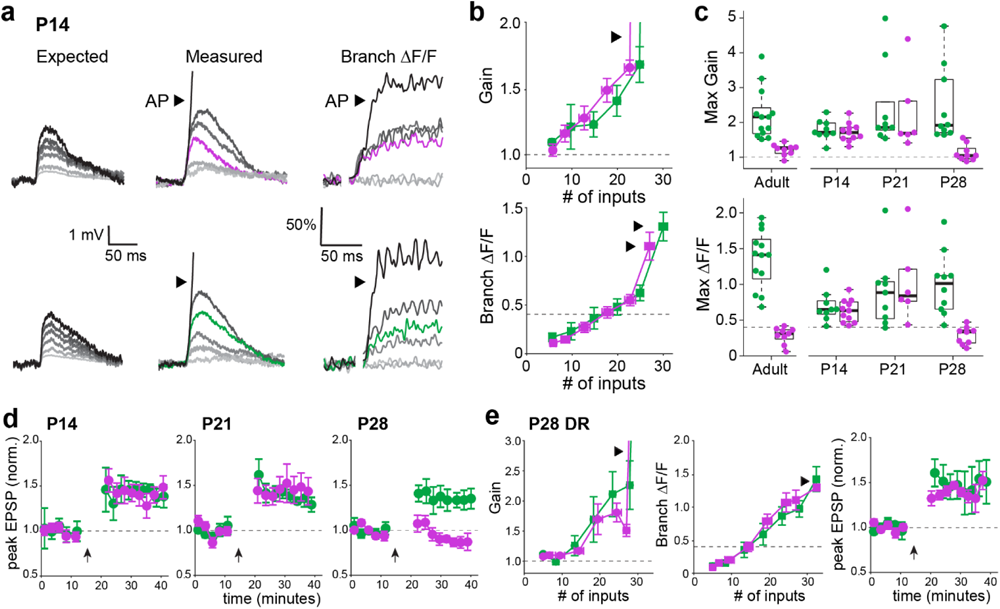
Oblique dendrites lose NMDAR-mediated properties after an experience-dependent critical period in postnatal development. a. Expected (left), measured (middle), and ΔF/F (right) for oblique (top) and basal (bottom) dendrites from the same neuron from a P14 mouse. Note supralinear integration and large Ca^2+^ signals in both branches. Both are driven to action potential initiation (AP, arrow). b. P14 population gain (measured/expected, left) and branch ΔF/F (right) (n = 9 basal branches and 9 oblique branches, both from 5 animals). Basal vs oblique gain*, p = 0.33; ΔF/F*, p = 0.70. c. Maximum gain (left) and local branch ΔF/F (prior to AP initiation, right) across developmental time points (Adult, P56+: n as in Fig. 1, basal vs oblique gain*, p = 1.07E-04; ΔF/F*, p = 1.07E-04. P14: n as in b, basal vs oblique gain*, p = 0.54; ΔF/F*, p = 0.76. P21: n = 9 basal branches, 4 animals, 6 oblique branches, 2 animals, basal vs oblique gain*, p = 0.52; ΔF/F*, p = 0.68. P28: n = 10 basal branches, 5 animals; 9 oblique branches, 5 animals, basal vs oblique gain*, p = 2.17E-05; ΔF/F*: p = 4.33E-05. Boxplot parameters as in Fig. 2. d. Plasticity (as described in Fig. 2) in basal versus oblique dendrites at P14 (n = 5 basal branches from 5 animals and 5 oblique branches from 5 animals, basal vs oblique post-induction*, p = 0.79), P21 (n= 5 basal branches from 4 animals and 7 oblique branches from 6 animals, basal vs oblique post-induction*, p = 0.87), and P28 (n = 7 basal branches, 5 animals; 7 oblique branches, 4 animals, basal vs oblique post-induction*, p = 1.25E-13). e. P28 population gain (measured/expected, left) and branch ΔF/F (middle) in dark-reared (DR) mice (n = 9 basal branches and 10 oblique branches from 4 animals, basal vs oblique gain*, p = 0.78; ΔF/F*, p = 0.96). Plasticity for dark-reared animals (right) (n = 5 apical branches from 4 animals and 6 oblique branches from 6 animals, basal vs oblique post-induction*, p = 0.81). *Mann-Whitney U test was used for all comparisons.

### Oblique dendrites develop distinct postsynaptic receptor composition

One possible interpretation of our results thus far is that adult oblique dendrites lack large local Ca^2+^ signals, integrate linearly, and do not potentiate due to a higher AMPA-to-NMDA ratio, as adult cortical AMPA receptors (AMPARs) are significantly less Ca^2+^-permeable than NMDARs^51,52^. To test this, we compared AMPAR and NMDAR-mediated responses at single spines from basal and oblique dendrites at P21 and in adults. Glutamate uncaging produced AMPAR-mediated EPSPs within the physiological range at single spines, below the threshold for spine Ca^2+^ influx (Fig. 4a, Extended Data Fig. 4). Mg^2+^-free ACSF containing the AMPAR antagonist DNQX was subsequently washed in and the uncaging protocol was repeated to assess synaptic NMDARs (Fig. 4b). The average maximum amplitudes of EPSPs under each condition were compared as AMPA/NMDA. At postnatal day 21, when oblique dendrites behave similarly to basal dendrites, spines from the two branch types had comparable AMPA/NMDA. By adulthood, this ratio increased for oblique dendrites (Fig. 4c), indicating relatively less NMDAR-mediated synaptic conductance, consistent with linear integration, smaller local Ca^2+^ signals, and a decreased capacity for synaptic potentiation.

**Figure 4:**
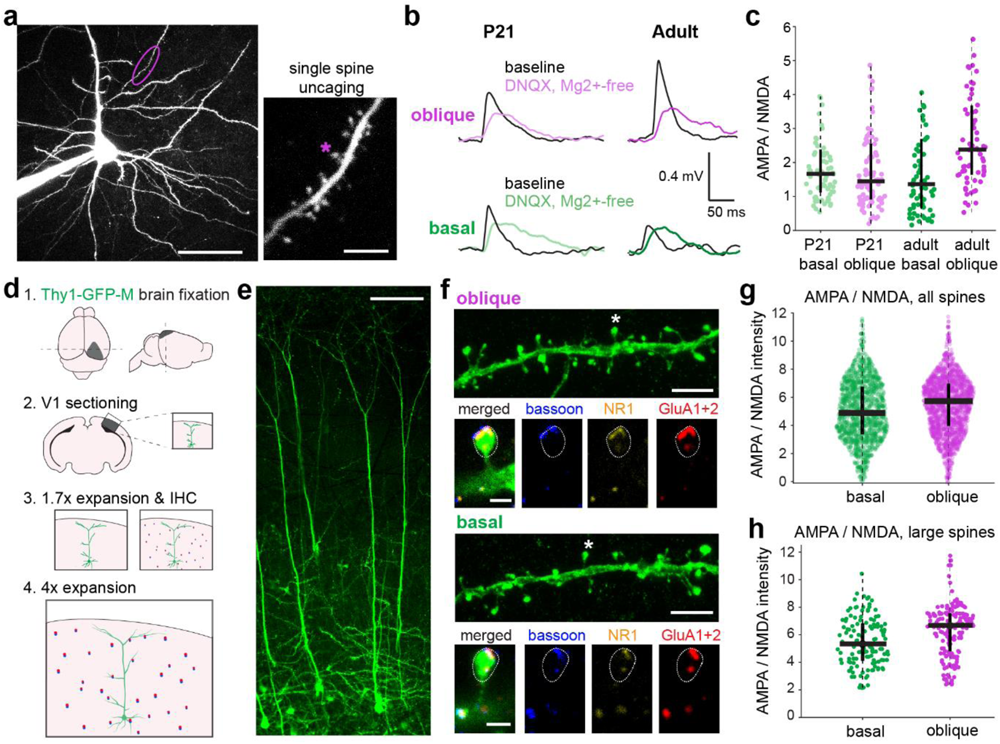
Changes in synaptic AMPA-to-NMDA ratio at oblique versus basal dendrites underlie differences in integration and plasticity. a. Example two-photon image of a V1 L5b PN from a P21 mouse (scale bar: 50 µm). An oblique dendrite (circled) is shown at higher magnification (right, scale bar: 5 µm). Uncaging site indicated by star. b. Representative average EPSPs from uncaging at single spines in P21 (left) or adult (right) mice from oblique (top) and basal (bottom) dendrites. Black traces: control, colored traces: the same spine in DNQX+Mg^2+^-free ACSF. c. Functional single spine AMPA/NMDA from basal and oblique dendrites in P21 and adult mice (P21: n = 72 basal spines from 4 mice and 89 oblique spines from 4 mice. Adult, P56+: n = 66 basal spines from 4 mice and 67 oblique spines from 5 mice). Outliers are not shown. P21 basal vs oblique, Mann-Whitney U test: p = 0.23; adult basal vs oblique, Mann-Whitney U test: p = 1.00E-04. Boxplot parameters as in Fig. 2. d. Schematic of the epitope-preserving Magnified Analysis of the Proteome (eMAP) experimental procedure. e. Confocal image of L5 PNs from a Thy1-GFP M P56+ mouse after 4x eMAP expansion. Scale bar: 300 µm. f. Example confocal images of expanded oblique (top) and basal (bottom) dendrites from the same neuron. For each branch, immunostaining for bassoon, NR1, and GluA1+2 is shown for one spine (indicated by star). Scale bar: 10 µm, inset: 2 µm. g. Ratio of AMPA/NMDA intensities for all spines: Mann-Whitney U test: p = 8.30E-11. h. Ratio of AMPA/NMDA intensities for the largest 10% of basal and oblique spines (n = 131 basal dendrite spines and 171 oblique dendrite spines). Mann-Whitney U test: p = 1.08E-04.

To provide complementary proteomic evidence for our observed changes in functional AMPA/NMDA, we acquired super-resolution images of oblique and basal dendrites and their synaptic receptor content using epitope-preserving Magnified Analysis of the Proteome (eMAP, Fig. 4d)^53,54^. Fixed slices from V1 of adult Thy1-GFP-M mice were expanded to 4x and stained with antibodies against the presynaptic marker bassoon, the obligate NMDAR subunit GluN1, and GluA1 and GluA2, the predominant AMPAR subunits in cortex^55^. Oblique and basal dendrites from L5b PNs were imaged with a confocal microscope in ∼40 µm-thick segments (∼180 µm expanded), and dendritic protrusions were annotated (Fig. 4e, f). Immature spines and protrusions without bassoon were excluded post hoc (although basal and oblique branches had similar numbers of immature and mature spines; Extended Data Fig. 4). For each spine head, the intensity of each fluorophore was spatially integrated (Fig. 4g), and the ratio of GluA1+GluA2/NR1 was taken. Consistent with our functional data, oblique dendrite synapses exhibited higher AMPA/NMDA than basal dendrites (Fig. 4h, Extended Data Fig. 4). This effect was pronounced in large spines (Fig. 4i). Taken together, these results indicate that oblique dendrites in adults have relatively less NMDAR content and conductance, and this underlies the differences in integration and plasticity rules between basal and oblique dendritic compartments.

### Compartmentalized synaptic stability and flexibility in single neurons in vivo

Variable levels of synaptic Ca^2+^ from NMDARs trigger molecular cascades that form, enlarge, shrink, and/or eliminate spiny synapses^56–62^. Given that we observed relatively low NMDAR conductance and expression in apical oblique dendrites, we hypothesized that oblique dendrites would have largely stable spiny synapses in vivo, whereas spiny synapses on basal dendrites should exhibit dynamic structural plasticity. To investigate this, we longitudinally imaged eGFP-expressing L5b PNs in adult mice. With extremely sparse cellular labeling and an optimized two-photon microscopy set up (Fig. 5a), we were able to resolve spines in oblique and basal branches in layers 4 and 5 (Fig. 5b). Across 5 or more consecutive days, we tracked spine dynamics for both basal and oblique dendritic compartments in individual cells (n = 6 cells, 775 basal spines and 794 oblique spines, 4 animals). Spine density did not significantly differ between basal and oblique compartments, consistent with our observations with eMAP (Fig. 5b, Extended Data Fig. 5). The overall survival fraction of basal and oblique spines was similar to what was previously reported for V1 apical tuft spines over a similar period of time, but spines on oblique branches had significantly higher survival rates (Extended Data Fig. 5, see ^63,64^). When considering both the formation and elimination of all spines between days (daily turnover ratio), oblique spines were remarkably stable, showing less than half the average turnover found on basal dendrites in the same cell (Fig. 5d). When individual spines were categorized by lifetime, where less than 4 days generally represents a “transient” synapse^64–68^, a larger fraction of basal dendritic spines had shorter lifetimes than oblique spines (Fig. 5a,e). Our results demonstrate that L5b PNs in adult mice possess a dendritic domain with highly stable synapses, in contrast to other dendritic domains with much higher rates of synaptic formation and elimination.

**Figure 5:**
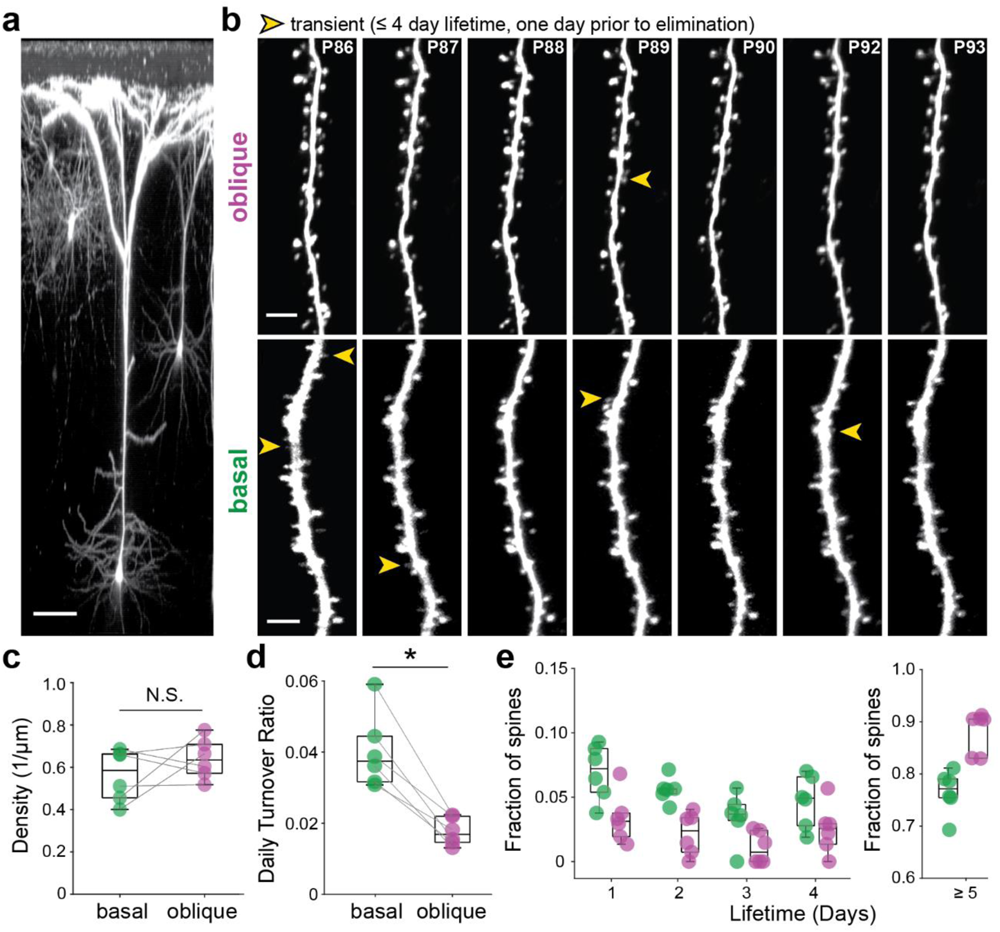
In vivo synaptic stability and flexibility at distinct dendritic compartments. a. 3D projection of an eGFP-expressing L5b PN, imaged in vivo with two-photon microscopy. Scale bar: 50 µm. b. Representative oblique (top) and basal (bottom) dendrites from the same layer 5 PN imaged across 7 days. All transient spines (with lifetimes of 4 days or fewer) are labeled (arrowheads) on the day prior to disappearance. Scale bar: 10 µm. c. Average spine density per cell for basal and oblique dendrites. Wilcoxon signed-rank, p = 0.56. d. Average daily turnover, per cell. Wilcoxon signed-rank, p = 0.03. e. Fraction of transient (≤ 4 day lifetime) and persistent (≥ 5 day lifetime) spines on oblique (purple) and basal (green) dendrites, per cell. Friedman’s test, p = 1.07E-06. For b-e, n = 6 cells with 775 basal spines and 794 oblique spines from 4 animals.

## Discussion

The plasticity of spiny synapses is widely considered to be the biological substrate of learning and memory. The current model posits that smaller dendritic protrusions are involved in the acquisition and refinement of information, whereas larger dendritic spines store acquired information^58,63,65,67–70^. The long-term stability of specific spiny synapses is not well understood and likely involves a combination of mechanisms, from network-level properties to molecular regulation. Here, we show that entire dendritic domains can be highly stable and resistant to long-term potentiation through a relative downregulation of synaptic NMDARs. These dendrites have comparable spine densities and morphologies to neighboring, plastic dendritic compartments but lack NMDAR-mediated supralinear integration and potentiation.

Our results do not rule out all possible plasticity mechanisms in oblique dendrites, but limited NMDAR-mediated Ca^2+^ influx constrains their potential to engage conventional molecular cascades^71^. In V1 L5b PNs, NMDAR-mediated properties decline in thalamorecipient dendrites after a critical period of early visual experience, a process which can be prevented by delaying the onset of vision. Thus, critical periods—the developmental window of experience required for normal connectivity and function^50,72,73^—occur at the subcellular level, an unprecedented degree of specificity, due to dendritic compartmentalization. Our findings establish a tractable new model for future studies on the formation and regulation of stable dendritic domains, which likely includes activity-dependent transcription, protein expression, and subcellular localization^74–78^.

Protecting existing knowledge while continually acquiring new information is a long-standing challenge for models of plasticity and biologically-inspired neural networks, which are prone to instability and catastrophic forgetting^1,5^. To resolve this plasticity-stability dilemma, some models decrease the plasticity of important weights^79–81^. Here we report the first biological evidence for these theoretical limitations: after an initial strengthening in early development, synapses on oblique dendritic spines are effectively stabilized through a domain-wide relative decline in NMDAR expression and function. Our results are distinct from—but may interact with—other models and observations that work to stabilize either plastic networks or individual synapses^21,82–84^. In primary visual cortex, the stable dendritic domain receives input from the visual thalamus, the definitive input to primary visual cortex. PNs in all layers of primary sensory cortices receive both first-order thalamic and intracortical inputs^38,40,85,86^, and this mechanism may help sensory neurons maintain fundamental sensory representations throughout a lifetime of learning and experience-dependent plasticity. Our results may also apply to other cortical areas: the oblique dendritic domain in mouse association cortex similarly lacks NMDAR-mediated supralinear integration^36^. Using this mechanism, cortical neurons may limit plasticity within specific dendrites to maintain essential, stable representations without compromising higher-order input plasticity. The restriction of plasticity within select dendritic compartments is a powerful mechanism by which single neurons can solve the tradeoff between flexibility and stability inherent to all learning systems.

## Acknowledgements

We thank Mark Bear, Elly Nedivi, Josh Trachtenberg, and the Harnett laboratory for their comments and suggestions and Kwanghun Chung for generously sharing equipment. This work was funded by a Life Sciences Research Foundation postdoctoral fellowship (C.E.Y.), and a Vallee Foundation Scholars Award, the James W. and Patricia T. Poitras Fund, and the National Institutes of Health R01NS106031 (M.T.H.).

## Author Contributions

C.E.Y. and M.T.H. conceived of the experiments. C.E.Y. performed patch-clamp physiology experiments. D.V. conducted eMAP experiments. N.J.B. performed sCRACM viral injections and histology. T.L.D.P performed viral injections and cranial window surgeries for in vivo two-photon imaging, and Q.Z. performed deep-layer spine imaging with C.E.Y, supervised by N.J. C.E.Y. carried out all analyses and prepared the figures. M.T.H. supervised all aspects of the project and wrote the manuscript with C.E.Y.

## Methods

### Animals

All animal procedures were carried out in accordance with NIH guidelines for animal research and were approved by the Massachusetts Institute of Technology Committee on Animal Care and the Animal Care and Use Committee at the University of California, Berkeley. C57BL6 male and female mice (Charles River Laboratories) were used in approximately equal numbers. Mice 8 weeks and older (P56+) were used for adult mouse experiments. For developmental timepoints, mice ages P10-P14, P18-22, and P28-P32 were used. All animals were kept in conventional social housing with unlimited food and water on a 12-hour light/dark cycle. For dark-rearing experiments, pregnant dams were moved to a dark room at E18 and pups were born and raised in complete darkness, with red light exposure during cage changes. Dark reared mice were ages P28-P32 at the time of experimentation. 8-week-old Thy-1-GFP-M mice (Jackson Laboratory) were used for eMAP experiments.

### Slice preparation

Sucrose-containing artificial cerebrospinal fluid (sACSF) was used during the slicing procedure, containing (in mM): 90 sucrose, 60 sodium chloride, 26.5 sodium bicarbonate, 2.75 potassium chloride, 1.25 sodium phosphate, 9 glucose, 1.1 calcium chloride, and 4.1 magnesium chloride, with an osmolality of 295-302. The sucrose solution was partially frozen to create a small amount of slush and was kept ice-cold. Artificial cerebrospinal fluid (ACSF) was used for recovery and recording, containing (in mM): 122 sodium chloride, 25 sodium bicarbonate, 3 potassium chloride, 1.25 sodium phosphate, 12 glucose, 1.2 calcium chloride, 1.2 magnesium chloride, 1 ascorbate, and 3 sodium pyruvate, with an osmolality of 302-307. All solutions were saturated with carbogen, 5% CO2 and 95% O2. Acute slice preparation was consistent for all ages and experiments. Mice were put under isoflurane-induced anesthesia and decapitated. The brain was extracted in ice-cold sucrose solution in less than one minute. The brain was blocked at a moderate angle (approximately 20 degrees from coronal) to favor the preservation of apical dendrites in the occipital cortex. After the brain was mounted and submerged in ice-cold sACSF, a vibratome (Lieca VT1200S) was used to cut 300 µm-thick slices, which were transferred to ACSF for 30-50 minutes at 36°C. Longer recovery times were used for tissue collected from younger animals. Following recovery, slices were kept at room temperature.

### Patch-clamp recording

Recordings were performed in ACSF (concentrations noted above) at 34–36 °C. Intracellular recording solution contained (in mM): 134 potassium gluconate, 6 KCl, 10 HEPES, 4 NaCl, 4 Mg2ATP, 0.3 NaGTP, and 14 phosphocreatine di(tris). Depending on the experiment, 0.05 mM Alexa 594, 0.1 mM Alexa 488, and/or 0.1 mM OGB-1 (Invitrogen) were added to the internal solution. An Olympus BX-61 microscope with Dodt optics and a water-immersion lens (60X, 0.9 NA; Olympus) was used to visualize cells. Whole-cell current-clamp recordings were obtained with a Dagan BVC-700A amplifier. Patch pipettes with thin-wall glass (1.5/1.0mm OD/ID, WPI) and resistances of 3-7 MΩ were used. Pipette capacitance was fully neutralized prior to break-in, and series resistance was kept fully balanced throughout, ranging from 5-30 MΩ. The liquid junction potential was not corrected. Current signals were digitized at 20 kHz and filtered at 10 kHz (Prairie View). L5b PNs were characterized by their large somas in layer 5, thick apical dendrites, broad arborization in layer 1, low input resistance, and prominent voltage sag.

### Virus injection

For sCRACM experiments, AAV2-hSyn-hChR2(H134R)-mCherry (UNC Vector Core) was expressed in neurons in the dorsal lateral geniculate nucleus. Mice 6 weeks or older were injected 4 weeks or more before slice preparation. Under isoflurane anesthesia, mice were secured on a stereotaxic apparatus with a feedback-controlled heating pad (DC Temperature Control System, FHC). Slow-release buprenorphine (1 mg/kg) was administered subcutaneously. After scalp incision, a small burr hole was drilled over the injection site at the following coordinates relative to bregma: anterior-posterior 2.7-2.8 mm; medio-lateral 2.3-2.4 mm. A beveled microinjection pipette containing was lowered to a depth of 2.7-2.9 mm, and approximately 100 nL of virus was injected at a rate of 50 nL/min using a Nanoject. Following virus injection and a five-minute rest, the pipette was removed and the incision was sutured. Accuracy of the injection was assessed in acute slices using two-photon microscopy. For image clarity, histology in Figure 1 was taken from a brain that was perfused, stored at 4°C overnight in 4% paraformaldehyde, transferred to PBS, and sectioned in 100 µm-thick slices. Sections were mounted, coverslipped, and imaged under a confocal microscope (Zeiss LSM 710 with a 10x objective, NA 0.45).

### sCRACM

After 4 weeks or more of ChR2 expression in dLGN, acute slices of V1 were prepared. A two-photon laser scanning system (Bruker), with dual galvanometers and a MaiTai DeepSee laser, was used to confirm the presence of mCherry-expressing cell bodies in the dLGN. For all recordings, TTX (1 µM) and 4-AP (100 µM) were added to ACSF to isolate monosynaptic connections. Following whole-cell configuration, a 5 ms 473 nm full-field LED was used to determine if the neuron received inputs from axons containing ChR2. If the LED drove EPSPs or spikes, a 9 by 17 (400 x 800 µm) stimulation grid with 50 µm spacing was positioned over the area containing the entire neuron, aligned at the pia using two-photon imaging under low magnification (4x, 0.16 NA air objective, Olympus) and Prairie View software. The stimulation grid controlled the location of a 473 nm laser beam (OptoEngine, LLC), and the duration of the light pulses was kept to 1-3 ms under <2 mW power. Each point on the grid was stimulated in order, progressing forward by column, with an interpoint delay of 1 s. 3-6 rounds of stimulation were averaged for each neuron. Baseline voltage was defined as the 50 ms before stimulation, and EPSP amplitudes were calculated as the baseline-subtracted maximum voltage within 50 ms after photostimulation. Population data was obtained by rotating the maps to align to the pia, overlaying the maps to align at the soma, normalizing peak voltages, and averaging across experiments.

### Glutamate uncaging

Following whole-cell dialysis with a structural dye via the patch pipette, basal or apical oblique branches were localized using two-photon imaging. A pipette containing caged glutamate (4-methoxy-7-nitroindolinyl-caged-L-glutamate or MNI-glutamate) diluted in ACSF (10 mM) was positioned just above the slice and over the recording site, and a picrospritzer (Picospritzer III, Parker Hannifin) was used to puff a constant flow of caged glutamate into the region of interest. A second Mai-Tai DeepSee laser controlled by Prairie View software was used for photostimulation of MNI-glutamate at 720 nm. Uncaging was targeted just adjacent to spine heads. In the imaging pathway, a linescan was used for simultaneous calcium imaging. Laser intensity for both lasers was independently controlled (Conoptics). Linescan imaging was performed at 1300 Hz, with a dwell time of 8 µs and a total scan time of less than 250 ms, with baseline fluorescence kept minimal and monitored throughout. Cells with signs of photodamage were excluded. The uncaging laser was calibrated to either a threshold calcium signal (∼100% ΔF/F) or an action potential, where the minimum power needed to drive either event was identified and used for all points. The uncaging dwell time was 0.2 ms, and the uncaging interval between multiple spines was 0.32 ms. Groups of 2-5 spines were stimulated independently and then in combination with other groups up to a total of 25-40 spines. Expected values were calculated by summing the average response of each group of spines. Ca^2+^ signals are expressed as (*F* − *F*_baseline_)/*F*_baseline_). For acute blockade of NMDA receptors, the competitive antagonist D-AP5 (50 µM) was used.

### Burst-timing dependent plasticity

For local branch stimulation, theta glass (2.0/1.4mm OD/ID) housing a bipolar stimulating electrode (ISO-Flex, AMPI) was positioned within 10 µm of a basal or apical oblique branch. Stimulation intensity was calibrated to generate a small EPSP of 1-3 mV, typically requiring 5-20 µA of current from the stimulating electrode. Ten minutes of baseline stimulation at 0.1 Hz was recorded to ensure stability of the EPSP. A -50 pA hyperpolarizing step was included to estimate input resistance. For plasticity induction, EPSPs were paired with a somatic current injection of 400-700 pA for 20 ms duration to produce a train of 2-4 action potentials. The peak of the first action potential was timed to be within 10 ms of the peak of the EPSP. Pairs were done in sets of 5 at 10 Hz, and sets of 5 were repeated 30 times at 0.1 Hz. After the induction period, EPSPs were monitored for at least 20 minutes. Inclusion criteria were as follows: resting membrane potential could fluctuate from baseline by no more than 3 mV, input resistance could not increase more than 20% of baseline; and series resistance had to be compensated fully throughout (<30 MΩ). Recordings were tested for baseline stability post-hoc using either correlation or linear regression (data not shown), and cells with significant changes in baseline recording were not included. The necessity of pre-post pairing was shown using the same recording set up but driving either the synaptic stimulation or the post-synaptic burst (and not both) during the induction period. For acute blockade of NMDARs, the competitive antagonist D-AP5 (50 µM) was used. Throughout plasticity experiments, picrotoxin (100 µM) was present in the bath to block GABAergic transmission in acute slices from adult mice and at P28. 10 µM picrotoxin was used for acute slices at P21, and no picrotoxin was needed at P14.

### Epitope-preserving Magnified Analysis of the Proteome

Mice were perfused with cold PBS followed by cold 4% PFA while under deep anesthesia (5% isoflurane). Brains were removed and kept in the same fixative overnight at 4°C and then washed with PBS at 4°C for at least 1 day. 1.0 mm coronal slices of primary visual cortex were cut on a vibratome and kept in PBS at 4°C until the day of processing. Slices were then incubated eMAP hydrogel monomer solution (30% acrylamide [A9099, MilliporeSigma, St. Louis, MO, USA], 10% sodium acrylate [408220, MilliporeSigma], 0.1% bis-acrylamide [161-0142, Bio-Rad Laboratories, Hercules, CA, USA], and 0.03% VA-044 (w/v) [Wako Chemicals, Richmond, VA, USA] in PBS]), protected from light, at 4°C overnight. For gelation, slices were mounted between glass slides in eMAP solution and sealed in a 50 mL conical tube with nitrogen gas at positive pressure of 10-12 psi at 37°C for 3 hours. The excess gel around the slices was then removed. To reach a first expansion stage of 1.7x, the slices were incubated overnight in a solution of 0.02% sodium azide (w/v) in PBS at 37°C. Slices were trimmed to contain only parts of primary visual cortex and further sectioned with a vibratome to 75 μm thickness (corresponding to ∼40 μm thickness of the pre-expanded tissue). Slices containing good candidate cells – L5 PNs whose apical trunk could be reconstructed at its full length in a single slice or at most two consecutive slices – were selected during live low-resolution confocal imaging sessions. These slices were trimmed to smallest possible samples of approximately 1.0 mm in both width and length. Slices were incubated in tissue clearing solution (6% SDS (w/v), 0.1 M phosphate buffer, 50 mM sodium sulfite, 0.02% sodium azide (w/v), pH 7.4) at 37°C for 6 hours, followed by incubation in preheated clearing solution at 95°C for 10 min. Cleared samples were thoroughly washed with PBS + 0.1% Triton X at 37 °C. Primary antibody staining was performed at 37°C overnight with the following antibodies: Anti-GFP (Life Technologies A10262), Anti-NMDAR1 (SYSY 114011), Anti-AMPAR1 (SYSY 182003), Anti-AMPAR2 (SYSY 182103), and Anti-Bassoon (SYSY 141004). For secondary staining, the following fluorescent antibodies were used: Bassoon: anti-Guinea pig-405 (AbCam ab175678); GFP: anti-Chicken-488 (Invitrogen A11039); NMDAR1: anti-Mouse-AF+555 (Invitrogen A32727); and AMPAR1 and AMPAR2: anti-Rabbit-AF+647 (Invitrogen A32733). Final expansion was performed just before imaging by putting the trimmed slices in 0.1 mM tris in distilled water, and approximately 4X total linear expansion was achieved. Slices were imaged using Leica TCS SP8 upright confocal DM6000 microscope equipped with a 63x HC PL APO CS2/1.2 W objective, hybrid detectors, and a white light laser. Within slices, intact L5b PNs were identified by their thick trunks and broad apical tufts. Within single neurons, both basal and oblique dendritic branches were imaged. To avoid photobleaching effects, no slice was imaged more than once, and basal and oblique dendrites were imaged in alternating order for each cell. Dendritic protrusions were analyzed using Fiji software. To draw regions of interest within spine heads in the GFP channel, a custom-written macro code was used to apply a median blur (2 pixels), threshold the image, and draw an ROI on the annotated protrusion. Using custom-written code in MATLAB, all color channels were thresholded to include only signals 2 S.D. above the mean fluorescence intensity, and signals from each channel were extracted from the ROI. The plane with peak antibody fluorescence was identified within the spine head ROI, and each channel’s fluorescence signals within the plane were summed. Long, thin dendritic protrusions without enlarged heads were classified as filopodia (head diameter/neck diameter < 1.3) and excluded from the final analysis, as were spines with no detectable bassoon.

### Ultrasparse eGFP expression in L5b PNs for deep layer structural imaging

Mice were approximately 8 weeks of age at the time of injection and cranial window surgery. Prior to surgery, mice were subcutaneously injected with dexamethasone (4mg kg^-1^) and buprenorphine (slow-release, 1 mg kg^-1^) and were anaesthetized with isoflurane (2% induction, 0.75-1.25% for maintenance). Mice were secured in a stereotaxic apparatus with a feedback-controlled heating pad. Eye ointment was applied to prevent dryness (Bepanthen, Bayer), and the scalp was shaved using hair removal cream and cleaned with iodine and ethanol. The skull was exposed, etched, and sealed with a light-curing glue (Adhese Universal, Ivoclar). A stainless steel headplate was secured to the skull with a light-curing flowable composite (Tetric EvoFlow Translucent, Ivoclar). Then a 3 mm-wide craniotomy was performed over V1 (2.5 mm lateral, 1 mm anterior to lambda). A 1:2 viral mixture of pENN.AAV.CamKII 0.4.Cre.SV40 (Addgene, diluted 1:100,000) and pAAV-hSyn-DIO-EGFP (Addgene) was injected at a volume of 150 nl at 3 sites, approximately 200–500 µm apart, at a depth of 550 µm. Following virus injection, the pipette was withdrawn after 5 minutes to avoid infecting superficial layers. Major blood vessels were avoided. Cranial windows consisted of a single 3 mm coverslip attached to a 4.5 mm x 2.5 mm coverslip (170 µm thickness, custom-made, Potomac). The outer ring of the cranial window was fixed to the skull with light-curing flowable composite (Tetric EvoFlow Translucent, Ivoclar). Following 1-2 weeks of recovery from surgery, animals were transferred to UC Berkeley and had 2 additional weeks of recovery.

*Longitudinal spine tracking in deep cortical layers with two-photon microscopy* Ultrasparse eGFP expression (1-2 cells per injection site) allowed us to visualize L5b PN dendritic spines 400-600 µm below the cortical surface with an optimized commercial two-photon microscope (Thorlabs Bergamo^®^ II Multiphoton). A femtosecond Ti:Sapphire laser (Coherent, Chameleon Ultra II) tuned to 920 nm was used as the excitation source for all experiments. A 25× water-immersion objective lens (Olympus, 1.05 NA, 2 mm WD) was used to focus the excitation light into the brain and collect the emitted fluorescence. Data acquisition and hardware were controlled using ThorImage software. Imaging started approximately 4 weeks after virus injection. Immediately prior to imaging, the mouse was anaesthetized (2% isoflurane) and then head-fixed under the microscope with continuous anesthesia (0.5–1% isoflurane). Body temperature was maintained using hand warmers. The headmount of the mouse was aligned so that its cranial window was perpendicular to the excitation beam. The correction collar of the objective lens was adjusted to minimize spherical aberration induced by the cranial window. L5b PNs were identified by cortical depth, with somas ∼500-600 µm below the pia, and by their thick apical dendrites and broad arborization in layer 1. For each cell, z-stacks were taken from the top-most oblique dendrite to the bottom-most basal dendrite (1 µm z steps, 203 µm x 203 µm field of view, 0.1 µm x/y pixel^-1^, 4 fps, 20 frames per step). The same cells and branches were imaged daily for 5-10 days. Post-objective power ranged from 38-65 mW for different cells, and care was taken to ensure that fluorescence levels were comparable throughout the tracking period.

### Image processing and spine tracking analysis

Two-photon z-stacks were analyzed in ImageJ (v1.54f). Smaller stacks of specific basal and oblique branches were sectioned from the original z-stack. Branches with steep changes in z were avoided. Each substack was rigidly aligned, gamma adjusted (1.5), and gaussian filtered (σ = 1). Branches from consecutive days were manually registered with a transparency tool (Vitrite), and similar lookup tables were used for each timepoint. Branches were only included if the dimmest protrusion was at least 5x higher than the standard deviation of the background around the dendrite. Dendritic protrusions were scored on maximum intensity z-projections and cross-checked with the original z-stack. The following criteria was used for scoring: protrusions must extend laterally, more than 0.5 µm from the branch. Protrusions that did not reach criteria at any point during the experiment were excluded from analysis. Daily tracking facilitated consistent identification of structures, even with small movements of the tissue. With this dataset, we calculated the fraction of protrusions that were present across the entire imaging period (survival fraction, SF) and the turnover ratio (TOR), defined as the fraction of structures that appear and/or disappear between days, as in ^89^ SF(t) = N(t)/N_0_, where N_0_ is the number of structures at t = 0, and N(t) is the number of structures of the original set surviving after time t. TOR (t_1_, t_2_) = (N_gained_+ N_lost_)/(N(t_1_) + N(t_2_)), where N(t_1_) and N(t_2_) are the total number of structures at the first and second time point. The lifetime of a given structure is the number of days present within the recording window.

## Data availability

All data and custom-written analysis code are available upon request.

