## Supplemental Figures for "A dendritic mechanism for balancing synaptic flexibility and stability"

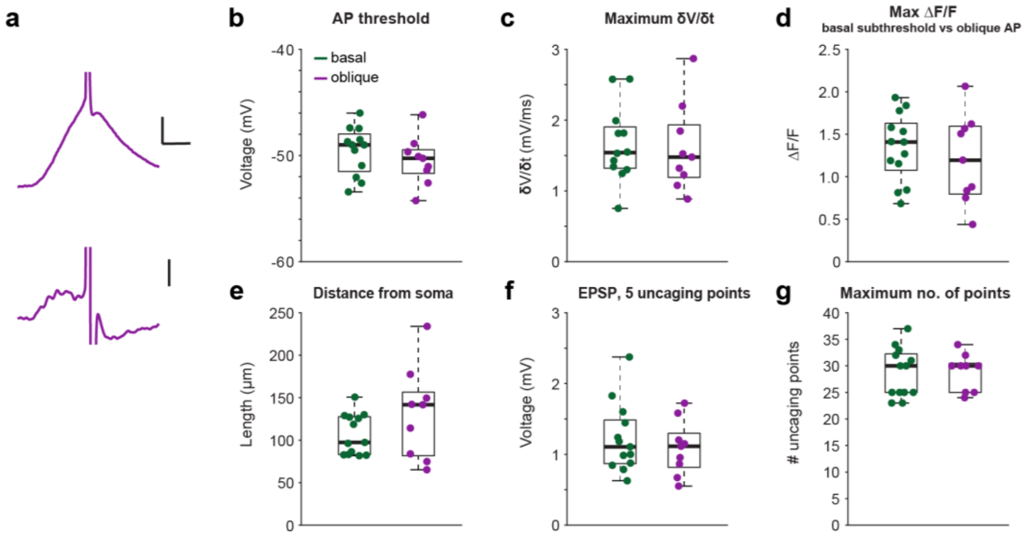


**Extended Data Figure 1. Properties of adult basal and oblique dendrites and experimental parameters of glutamate uncaging experiments.**

a. Example voltage waveform and corresponding δV/δt for an AP evoked by glutamate uncaging at an oblique branch. Scale bar, top: 5 mV, 10 ms, bottom: 1 mV/s.
b. Voltage thresholds for APs evoked by uncaging at basal (green) and oblique (purple) dendrites (n = 12 basal branches, 7 animals; 9 oblique branches, 6 animals. Mann-Whitney U test: $p = 0.27$).
c. Maximum δV/δt prior to AP initiation for uncaging at basal and oblique branches (n = 12 basal branches, 7 animals; 9 oblique branches, 6 animals. Mann-Whitney U test: $p = 0.64$).
d. Maximum ΔF/F for subthreshold responses at basal dendrites and maximum ΔF/F in oblique dendrites following an AP (n = 13 basal branches, 7 animals; 8 oblique branches, 6 animals. Mann-Whitney U test: $p = 0.69$).
e. Uncaging site distance from soma for basal and oblique dendrites (n = 13 basal branches, 7 animals, 9 oblique branches, 6 animals. Mann-Whitney U test: $p = 0.34$).
f. EPSP amplitude for basal and oblique dendrites driven by 5 uncaging points (n = 13 basal branches, 7 animals; 9 oblique branches, 6 animals. Mann-Whitney U test: $p = 0.64$).
g. Maximum number of uncaging points used in basal or oblique experiments (n = 13 basal branches, 7 animals; 9 oblique branches, 6 animals. Mann-Whitney U test: $p = 1$).

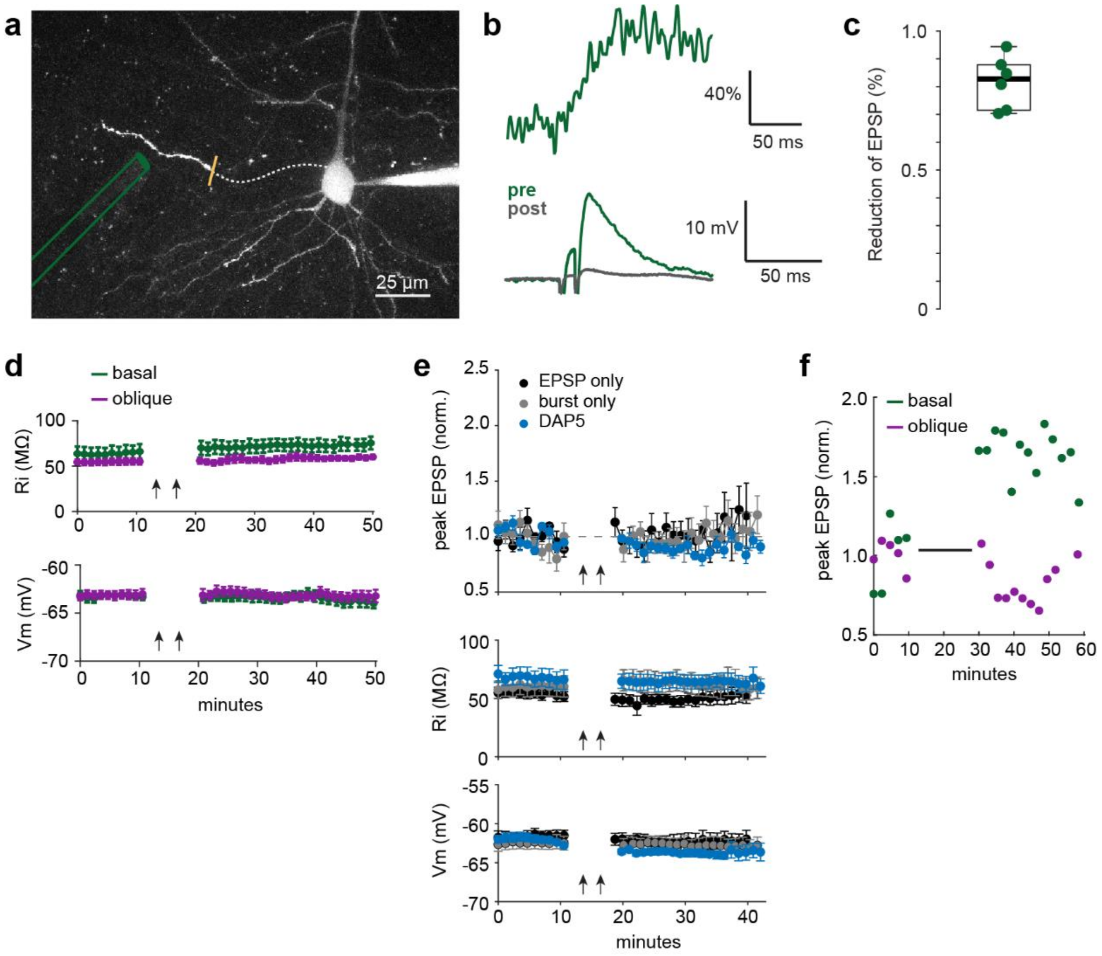


**Extended Data Figure 2. Experimental measurement of local electrical stimulation spatial spread and properties of synaptic potentiation in adult L5 PNs.**

a. Two-photon z-stack showing experimental set up. Theta glass pipette housing a bipolar electrode is positioned close to a basal branch to produce EPSPs and local $Ca^{2+}$ signals. The branch is then severed with a laser (yellow) and EPSPs are measured again. The damaged proximal part of branch is indicated with a dotted line for clarity.

b. A local branch $Ca^{2+}$ signal (top) and somatically-recorded EPSP (bottom) were driven in the basal branch shown in a, prior to the laser cut (pre, green). Following severing of the branch, the EPSP is greatly reduced (post, gray).

c. Percent reduction of peak EPSP amplitude following laser cutting, n = 6 basal branches from 6 neurons and 5 mice.

d. Input resistance (top) and resting membrane potential (bottom) monitored throughout adult plasticity experiments shown in Figure 2. Neither corresponded with changes in synaptic potentiation.

e. Coordinated pre- and post-synaptic activity and NMDA receptors are required for synaptic potentiation in basal dendrites. EPSPs (top) in basal dendrites following induction protocols with presynaptic-only stimulation (black, n = 6 cells, 5 animals), postsynaptic-only (gray, n = 7 cells, 4 animals), and pre/post pairing in the presence of D-AP5 (blue, n = 5 cells, 5 animals) with corresponding input resistance (middle) and resting membrane potential (bottom). Presynaptic-only or postsynaptic-only before versus after pairing comparisons were not statistically significant (pre-only, Mann-Whitney U test: p = 0.42, post-only, Mann-

Whitney U test: p = 0.66,); pairing with D-AP5 showed small but statistically significant depression, Mann-Whitney U test: p = 0.006.

f. EPSPs (2 min binned averages) from a basal and oblique dendrite before and after plasticity induction with 60 pairings. Despite double the number of pairings, the oblique branch does not potentiate.

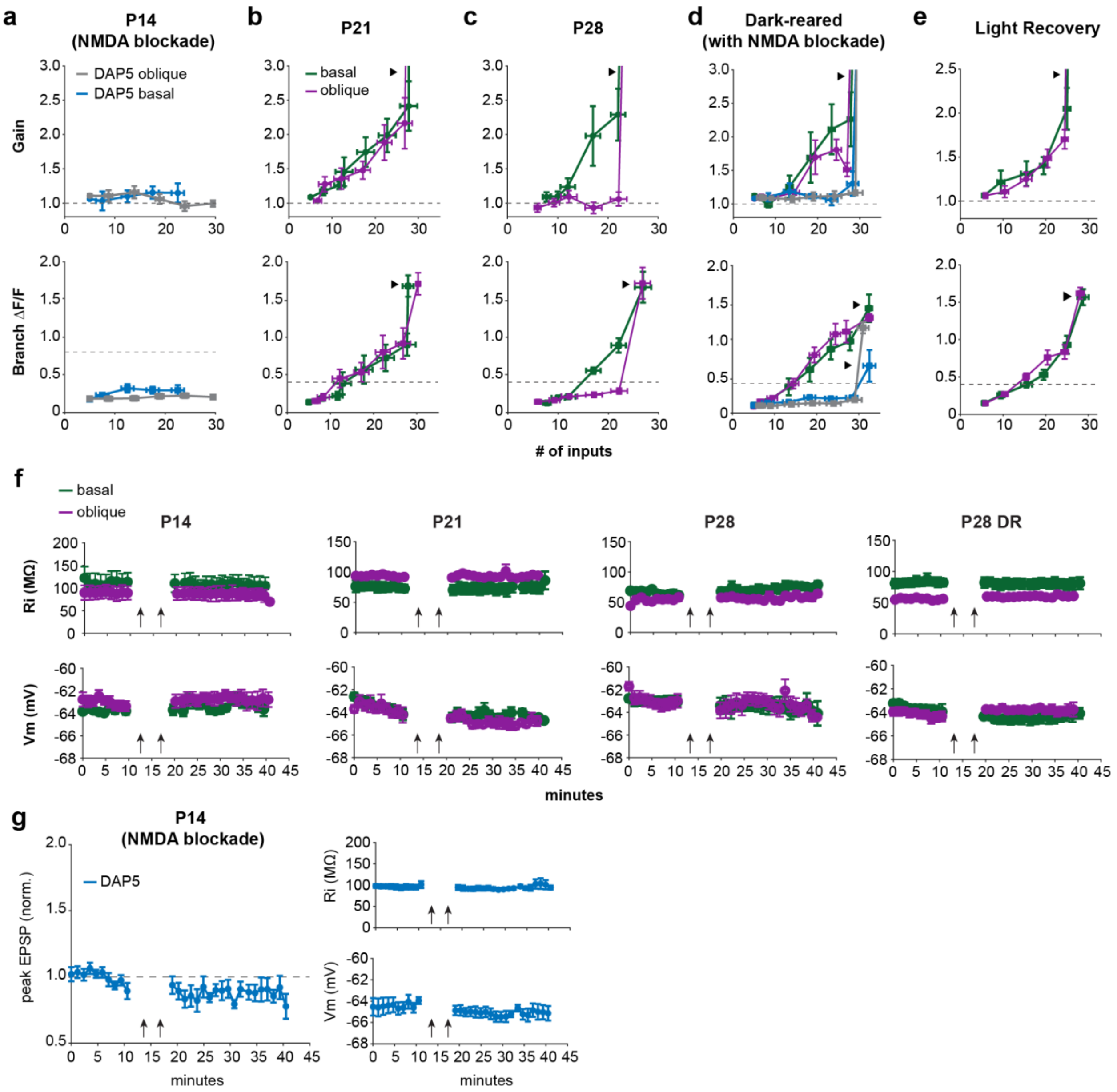


**Extended Data Figure 3. Dendritic input-output relationships and plasticity properties for developmental timepoints and controls.**

All panels are gain (top) and branch ΔF/F (bottom) vs. number of glutamate uncaging inputs.

a. Blockade of NMDARs by D-AP5 eliminates supralinear gain and branch calcium influx for both basal (blue) and oblique (grey) dendrites in P14 mice.
b. Basal and oblique dendrites exhibit supralinear gain and large $Ca^{2+}$ signals in P21 mice under control conditions. Basal vs oblique gain*: $p = 0.29$; ΔF/F*: $p = 0.49$.
c. In P28 mice, basal dendrites exhibit supralinear integration and large $Ca^{2+}$ signals, in contrast to oblique dendrites, which are linear and lack $Ca^{2+}$ signals prior to AP initiation. Basal vs oblique gain*: $p = 3.14E-06$; ΔF/F*: $p = 7.84E-04$. AP excluded in both comparisons.

d. Both basal and oblique dendrites in P28 dark-reared animals integrate supralinearly with large $Ca^{2+}$ signals. Basal vs oblique gain*: $p = 0.78$; ΔF/F*: $p = 0.96$, Mann-Whitney U test. These processes are NMDAR-dependent (D-AP5 blockade: basal, blue, oblique, grey).

e. Both basal and oblique dendrites in dark-reared animals that are returned to normal light conditions for 2-4 weeks retain supralinear integration and large $Ca^{2+}$ signals. Basal vs oblique gain*: $p = 0.75$; ΔF/F*: $p = 0.65$.

f. Corresponding input resistance (top) and resting membrane potential (bottom) during plasticity experiments in Fig 3, for each developmental timepoint and dark-reared animals.

g. D-AP5 prevents synaptic potentiation in P14 basal and oblique dendrites (n = 4 oblique dendrites in 3 animals, 2 basal dendrites in 2 animals, pre vs post induction*: p = 1.41E-06).

*Mann-Whitney U test was used for all comparisons.

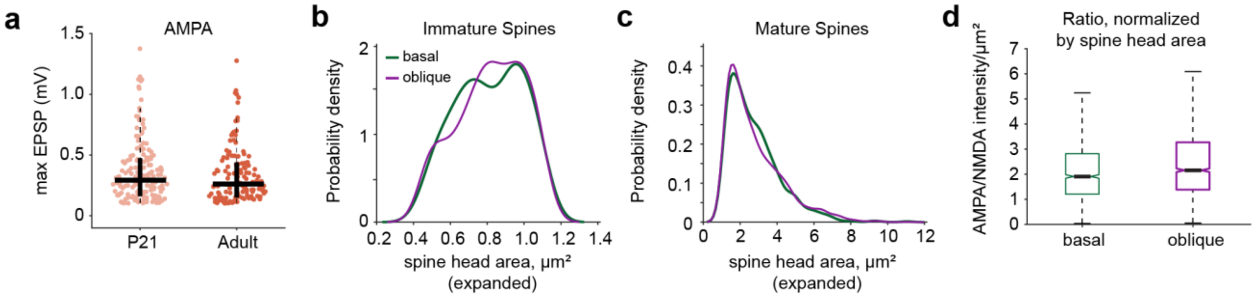


**Extended Data Figure 4. Amplitudes of uncaging-evoked single spine EPSPs and properties of spines in basal and oblique dendrites in expanded tissue.**

a. AMPAR-mediated EPSP amplitudes for all spines used to measure functional AMPA/NMDA in P21 and adult mice (Mann-Whitney U test: $p = 0.43$).
b. Probability density of immature spines in basal and oblique dendrites (n = 392 immature spines in basal dendrites, 513 immature spines in oblique dendrites, 13 cells, 3 animals, Mann-Whitney U test: $p = 0.34$).
c. Probability density of mature spines in basal and oblique dendrites (n = 1305 spines in basal dendrites, 1719 spines in oblique dendrites, 13 cells, 3 animals, Mann-Whitney U test: $p = 0.77$).
d. Ratio of AMPA/NMDA fluorescence intensity normalized by spine head area for spines in basal and oblique dendrites, Mann-Whitney U test: $p = 2.79E-08$.

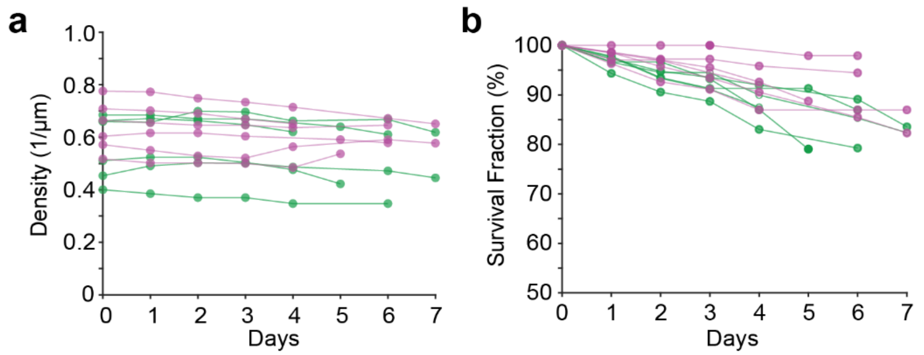


**Extended Data Figure 5. Spine tracking metrics across imaging days.**

a. Spine density for oblique (purple) and basal (green) dendrites, per cell. Daily basal vs oblique spine density*: Wilcoxon signed-rank, $p = 0.07$.
b. Percentage of oblique and basal protrusions surviving as a function of time, per cell. Daily basal vs oblique survival fraction*: Wilcoxon signed-rank, $p = 8.63E-04$.
* n = 6 cells with 775 basal spines and 794 oblique spines from 4 animals.
